# Root endophytes improve physiological performance and yield in crops under salt stress by up-regulating the foliar sodium concentration

**DOI:** 10.1101/435032

**Authors:** Marco A. Molina-Montenegro, Ian S. Acuña-Rodríguez, Cristian Torres-Díaz, Pedro E. Gundel

## Abstract

Increase in the soil salinity will be a conspicuous constraint for both native plant communities as well as several crops worldwide. In this context plant root-associated fungi appear as a new strategy to improve ecophysiological performance and yield of crops under abiotic stress. Here, we evaluated how the inoculation of fungal endophytes isolated from Antarctic plants can improve the ecophysiological performance in cultivars of tomato and lettuce, grown under different salt conditions. In addition, we assessed if the expression of the *NHX1* gene that is related with salt tolerance is enhanced in presence of fungal endophytes. Finally, we evaluated the regulation the nutritional status and specifically the Na^+^ content in leaves. Overall, those individuals with presence of endophytes showed higher ecophysiological performance. In addition, the presence of fungal endophytes was correlated with a higher regulation of ion homeostasis by enhanced expression of *NHX1* gene. Our results suggest that presence of fungal endophytes could minimize the negative effect of salt by improving osmotic tolerance through ecophysiological and molecular mechanisms. Thus, root-endophytes might be a successful biotechnological tool to maintain high levels of ecophysiological performance and productivity in zones under osmotic stress, acting as potential solution to maintain the global food security.

**Highlight:** We showed that functional symbiosis improve the physiological performance and yield in crops subjected to salinity, by biochemical and molecular mechanisms. Thus, here we pointed a successful strategy to meet the future challenges for help to maintain the food security.

## Introduction

The earth planet is facing dramatic environmental changes driven by the market economy with an increasing human population and higher life expectancy (Tilman et al., 2002; Foley et al., 2005; Godfray et al., 2010). Hence, food production may be compromised by a combination of effects from the undergoing climate change, land degradation and contamination. Although traditional breeding and biotechnology are likely to overcome part of these constraints by accommodating plant phenotypes for matching the environment (Carroll et al., 2014), a search for more ecological and friendly practices is a priority as a mean to reduce the use of agro-pesticides and/or enhance the environmental tolerance in agroecosystems. In this context, microbial symbionts of plants appears as a promising alternative for improving plant performance and maintain, or even increase, the yield of crops (Gundel et al., 2013; Kauppinen et al., 2016; Molina-Montenegro et al., 2016a; Hawkes and Connor, 2017; Wei and Jousset, 2017).

Soil salinization can be a symptom of land degradation, as a result of inappropriate cultural practices and excessive agricultural use, and it is currently affecting a vast territory of productive areas of the world (Metternicht and Zinck, 2003; Rengasamy, 2006; Qadir et al., 2014; Roy et al., 2014). As a consequence, plant resistance to salt, mainly to the sodium cation (Na^+^), is a desirable trait to be selected in cultivated plants. A mechanism of plant tolerance in glycophytic (i.e. salt susceptible) species consists of reducing the cytoplasmic sodium toxicity by pumping Na^+^ up into the vacuole through the tonoplast Na^+^/H^+^ antiporters (Blumwald et al., 2000; Munns and Tester, 2008). Apart from reducing sodium toxicity, increased ion concentration in vacuoles allows plants to alleviate water deficit and maintain positive carbon gain (Pardo and Quintero, 2002; Munns and Tester, 2008). This is governed by a family of genes *NHX*, well described in *Arabidopsis thaliana* but also present in other species such as tomato (Pardo et al., 2006). Tomato plants are able to accumulate sodium in leaves when growing in soils with high sodium contents (Zhang and Blumwald, 2001) and an increased tolerance has been observed in plants overexpressing *NHX* genes (Gouiaa and Khoudi, 2015; but see also Leidi et al., 2010). It has been also scrutinized on how plant tolerance to sodium can be improved by the association with root microorganisms (reviewed in Hanin et al., 2016; Rho et al., 2018). For example, inoculation with arbuscular mycorrhizal fungi (AMF) significantly improved yield in tomato under high salt content (Al-Karaki, 2000; Latef and Chaoxing, 2011). The concentration of Na was lower in AMF plants than in non-AMF plants in both, saline and no-saline soils, with the former ones displaying a higher activity of the enzymatic antioxidant machinery (Latef and Chaoxing, 2011). Taken all together, root symbiotic microorganisms seem to activate the immune system in plants increasing the overall performance, even under saline soils. Interestingly, it is unknown the link between the presence of (a) root symbiont(s) and a given specific molecular mechanism with physiological consequences by which a reduced toxicity to Na^+^ is observed in plants.

Accumulating evidence confirm that the association of plants with multiple microorganisms (e.g., fungal endophytes) either in roots or aboveground tissues is not the exception but the rule (Rodriguez et al., 2009; Hardoim et al., 2015; Jin et al., 2017; Lemanceau et al., 2017). Some of the oft-cited effects of fungal endophytes are host acquisition of new metabolic capabilities, source of secondary metabolites, and elicitation of plant defense systems against abiotic stress factors and enemies (Rodriguez and Redman, 2005; Pineda et al., 2010; Hamilton et al., 2012; Pieterse et al., 2014; Hawkes and Connor, 2017). Some aspects are today challenging when addressing the effects of root fungal endophytes on plant performance. First, while some studies focuses on soil microbiota as a whole (Vandenkoornhuyse et al., 2015), others try to identify the responsible microorganism of the main effect on the host plant fitness (Gallardo-Cerda et al., 2018). Second, although results clearly show positive effects on host plants at the individual level (i.e. yield or biomass increase), a key point is to unveil the molecular pathways and physiological effects of symbionts that underlie the host improvement under stress conditions. For example, tomato plants (*Solanum lycopersicum*) that were grown in native soils showed an elicited response in their immune systems related to protection against oxidative stress compared with their conspecific plants but grown in sterile soils (Chialva et al., 2018). This latter effect was partially reverted when the plants growing in sterile soils were inoculated with the AMF *Funneliformis mosseae* (Chialva et al., 2018), indicating a key role for that kind of root symbiont. Arbuscular mycorrhizal fungi also improved the general performance of *Robinia pseudoacacia* seedlings under stress by NaCl salt (Chen et al., 2017). The overall physiological status of the plants (photosynthetic rate, photosystem II quantum efficiency and relative water content) was boosted by the symbiont, at the time that genes encoding membrane transport proteins involved in K^+^/Na^+^ homeostasis in roots were found to be upregulated (Chen et al., 2017). Although AMF is by far one of the soil microorganism groups most studied likely due to its importance as provider mutualist (Omacini et al., 2012; Pieterse et al., 2014), other root colonizers (e.g., *Trichoderma* spp., *Piriformospora indica*, *Penicillium* spp.) are known to deliver benefits to their host plants. Strains of *Trichoderma*, a common soil fungus, increased the tolerance to Na+ in *Arabidopsis thaliana* by increasing the level of the auxin IAA (indole-3-acetic acid), production of osmolites and antioxidants (Contreras-Cornejo et al., 2014).

It has been proposed that symbiotic fungi are responsible of the adaptation of plants to environmental stress factors (Clay and Schardl, 2002; Redman et al., 2002; Pieterse et al., 2014). More recently, different bio-prospective research programs are considering fungi present in unexplored harsh environments, as an interesting strategy to finding new metabolic pathways and ultimately, new bioactive constituents (Godinho et al., 2013). In this way, Antarctica has truly unique ecosystems representing one of the most severe climatic conditions for life on earth; i.e., low water availability, high UV-B radiation, extreme low temperatures and saline soils (Convey et al., 2014). Nevertheless, there are plants and microorganisms that have evolved to inhabit in these harsh conditions, and in some cases these microorganisms have been found to be responsible of improving plant adaptation to such stressful conditions by establishing the so-called ‘functional symbiosis’ (see Torres-Diaz et al., 2016). Indeed, using antarctic plant species as a source, we recently identified from *Colobanthus quitensis* two species of *Penicilium* that succeed in retain their functional role observed in their native plant host when exposed to a new plant species ((Molina-Montenegro et al., 2016a). Thus, taking into account the geographic isolation and inhospitable conditions for growth that prevails in Antarctica and the referred ecological role of some antarctic plant endophytes; it is highly plausible to consider that these microorganisms may have unique metabolic pathways to produce or activate novel molecules. Since this display of interesting bioactivities highlights their potential as biotechnological tools; Antarctic endophytes should be suitable sources to assess the role of microbial plant symbionts in the adaptation / mitigation of crops under large environmental constraints, as suggested by climate change (IPCC, 2016).

In this article, we explored the effect of fungal endophytes isolated from roots of plants native to the Antarctic continent on the individual performance and final yield of two horticulture crop species, *Lactuca sativa* and *Solanum lycopersicum*, under stress by sodium chloride salt (NaCl). Our hypothesis was that the symbiosis with the root fungal endophytes benefits host plants but this effect is far more evident under stress condition. On the other hand, this endophyte-mediated improvement of host plant performance correlates with physiological processes at individual level such as photosynthesis and water use efficiency as well as with an upregulation of the gene involved in the vacuole sequestration of Na^+^. Our work gives insights on the underlying mechanisms by which beneficial fungal endophytes are associated with host plant benefits, and establish the foundations for the design of ecologically friendly practices by using of microorganisms in agriculture.

## Methodology

### Inoculum of fungal endophyte

We used two Antarctic fungal endophytes (AFE) isolated from the roots of two Antarctic plants, *Colobanthus quitensis* (AFE001) and *Deschampsia antarctica* (AFE002), during the growing season 2015-2016. Details concerning to the procedures of fungal isolations, molecular characterization and species identification of these two fungal endophytes can be found in a previous study (Molina-Montenegro et al., 2016a). The isolates AFE001 (Genebank accession number: KJ881370) and AFE002 (Genebank accession number: KJ881371) were identified as *Penicillium brevicompactum* and *P. chrysogenum*, respectively (Molina-Montenegro et al., 2016a). These inoculums are maintained as part of the collection of microorganisms of the Plant Ecology Laboratory, Universidad de Talca, Chile. The inoculums were separated in different Petri dishes and then frozen until to be used in the experiments.

Fresh inoculums were obtained during March 2017 from single-conidia of AFE001 and AFE002 cultured on potato dextrose agar (PDA) medium diluted eight times and supplemented with 50 mg/ml of streptomycin. Cultures with endophytes were incubated at 22 ± 2 °C with a photoperiod 14/10 day/night. After two weeks of incubation, conidia were harvested from plates by adding 10 ml of sterile water and gently scraping off conidia with a sterile glass slide. The conidia suspension was adjusted to 100 ml of 0.05% Tween-100, sterilized solution, filtered through three layers of sterile cotton cheesecloth gauze. Conidia concentration was estimated by using a Neubauer chamber and adjusted to 1 × 10 ^5^ conidia/ml and its viability was tested according to methodology described by Greenfield et al., (2016) and the mean conidia viability was > 95%.

### Host plant species and experimental design

*Lactuca sativa* (lettuce var. Romaine) and *Solanum lycopersicum* (tomato var. Moneymaker) seedlings were obtained from seeds germinated in glasshouse located at the Universidad de Talca, Talca, Chile (35.4° S), under semi-controlled environmental conditions of light and temperature (760 ± 96 μmol m^-2^s^-1^; 22 ± 5 °C, respectively). For treatment setup, lettuce and tomato seedlings were transplanted into the field when individuals presented at least four expanded leaves and 3-cm roots. One-hundred seedlings of each species were randomly assigned to one of the four treatments: (*i*) plants irrigated with 40 cc of tap water/day plus 50 mM NaCl solution, (*ii*) plants irrigated with 40 cc of tap water/day plus 50 mM NaCl solution and the presence of endophytes, (*iii*) plants irrigated with 40 cc of tap water/day plus 150 mM NaCl solution, and (*iv*) plants irrigated with 40 cc of tap water/day plus 150 mM NaCl solution and the presence of endophytes.

The endophyte inoculum consisted of a concentrated mix of conidia from the two fungi (*P. brevicompactum* and *P. chrysogenum*). The plant inoculation was repeated to ensure the fungi to establish an effective association; verification of the symbiosis was evidenced by microscopy re-cultured technics in plates. Before the beginning of the experiment, two plants of each species/treatment were sacrificed to check microscopically for the presence and/or absence of endophytes by routine staining of roots.

The amount of tap water that is normally added to lettuce and tomato crops in different commercial stations in the Maule region of Chile for this time of the year, ranges from 30 to 45 cc/day per plant. The seedlings (*n* = 100 for each treatment) were transplanted to the field and distributed in rows; distance between rows 0.5 m and distance between plants 0.2 m. Each treatment was assigned to independent rows with four rows per each treatment. The soil of the plot is characterized by high content of clay, good drainage, and low levels of salt and macronutrients. Each individual was fertilized with 0.2 g L^-1^ of Phostrogen^®^ (Solaris, NPK, 14:10:27) every 30 days. The experiment lasted for 90 (lettuce) and 100 (tomato) days, and the measurements were made simultaneously in all treatments for both species. Environmental conditions were recorded at midday (12:00-15:00h) during the whole experimental period. Air temperature and relative humidity was recorded with a data logger (HOBO-Pro v2 U-23) and sunlight was registered with a portable photosynthetic active radiation sensor (Li-190 quantum sensor). During the experiment, daily mean temperature and relative humidity were 21.6 °C (± 3.8) and 65 % (± 12), respectively; while daily mean radiation was 1.422 μmol m^-2^s^-1^ (± 336).

### Ecophysiological traits

The net photosynthesis rate (A), and transpiration rate (E) were measured on a visually healthy leaf from 25 individuals corresponding to each treatment of the factorial design. Measurements were made on the same individual at the day 30, 60 and 90 (lettuce) or 100 (tomato), by an infrared gas analyzer (IRGA, Infra Red Gas Analyser, CIRAS-2, PP-Systems Haverhill, USA). From gas exchange measurements, we estimated the instantaneous water use efficiency (WUE) for photosynthesis as the ratio between photosynthetic rate and transpiration (A/E). This parameter is an indicator of plant water stress in a microsite or condition, because an increase in WUE is usually induced by a decrease in water availability (Lambers et al., 1998).

### Crop yield

At the end of the experimental period, sampled individuals in each treatment were extracted from the soil without damaging the root system. Subsequently, the roots were washed without removing them from the stem and left to dry in the shade for 1 h. Total fresh biomass of both, shoots and roots of each individual was weighed with a digital electronic scale (Boeco BBL-52; 0.01 g-precision). Taking into account for each species the nature of their commercialized product, the individual average final crop yield was estimated after over-drying at 62 °C for 96 h the complete shoot tissues of each lettuce (i.e. the leaves), and the fruits in the case of tomato plants.

### Gene expression

Total RNA was extracted from leaves of 0, 30 and 90 (lettuce) or 100 (tomato) days old plants (*n* = 5 ind. per species-treatment) according to Chang et al., (1993). RNA yield and purity were checked by means of UV absorption spectra, whereas RNA integrity was determined by electrophoresis on agarose gel. DNA was removed using TURBO DNA-free (Applied Biosystems, California, USA) from aliquots of total RNA. The first strand cDNA was synthesized according to previous methods (Ruiz-Carrasco et al. 2011). The reaction of quantitative PCR (qPCR) was performed in a final volume of 20 μl containing the cDNA, 5 pmol of each primer and 12.5 μl of the Fast SYBR Green PCR master mix (Applied Biosystems) according to the manufacturer’s instructions. The Elongation Factor 1a (EF1a) housekeeping gene was used as reference gene to normalize, and estimate up- or down-regulation of the target genes for all qPCR analyses: 50-GTACGCATGGGTGCTTGACAAACTC-30 (forward); 50-ATCAGCCTGGGAGGTACCAGTAAT-30 (reverse). NHX1 sequences were used to amplify LsNHX1 amplicons that were 200-bp long: 50-GCACTTCTGTTGCTGTGAGTTCCA-30 (forward); 50-TGTGCCCTGACCTCGTAAACTGAT-30 (reverse). PCRs were carried out with Step One Plus 7500 Fast (Applied Biosystems) for an initial cycle of 30min at 45° C and 2 min at 95° C, and then 40 cycles as follows: 95° C for 30 s, 60° C for 30 s, 72° C for 2min and finally one cycle at 72° C for 10 min. Cycle threshold (Ct) values were obtained and analyzed with the 2^ΔΔC^_T_ method (Livak & Schmittgen, 2001). The relative expression ratio (log_2_) between each target gene and the reference gene, and fold changes (FC) between drought-treated samples vs. corresponding controls were calculated from the qRT-PCR efficiencies and the crossing point deviation using the mathematical model proposed by Pfaffl (2001).

### Nutrient content

At the end of the experiment, the concentrations of elements (N, P, K and Na) and molecules (NO_3_^−^ and NH_4_^+^) in the plant produced biomass was determined. Determinations were done on seven individuals from each treatment, and expressed as percentage on dry weight basis. All analyse were conducted in the Laboratory of Nutrient Analysis at Universidad de Talca, Talca. Shoot nutrient concentrations were determined after dry-ashing (except for nitrogen). NO_3_^−^ and NH_4_^+^ were determinated after KCl extraction; P by Bray-1 method; K, and Na after ammonium acetate extraction. N was determined via combustion analysis (CNS-2000 Macro Analyzer, Leco Inc., MI, USA). P, K and Na were measured by ICP-OES (Perkin Elmer Optima 3000DV, Wellesley, MA, USA).

### Statistical analysis

Six response variables related with plant fitness, physiological performance, gene expression and nutritional status were analyzed to describe the role of the endophytes on the biological performance of lettuce and tomato under saline stress. A standard two-way analysis of variance (ANOVA) was used to evaluate the effect of saline stress and endophyte inoculation for the final yield and final Na+ foliar content. To evaluate the effect of saline stress and endophyte inoculation on those variables measured along time (photosynthesis, water use efficiency (WUE) and gene expression), we used two-way repeated measures (rm) ANOVAs. Model fitting was performed with the *aov* function from the base R options using the individual nested in time as the random error structure as allowed for homoscedastic and orthogonal designs. Shoot nutrient concentrations (N, P and K) under the four experimental treatments, control/E−, control/E+, salt added/E−, and salt added/E+, were compared for lettuce and tomato using a one-way ANOVAs. For the one-way and two-way standard ANOVA’s, *a posteriori* differences between treatments were evaluated using Honest Significant Differences (HSD) Tukey tests. Normality and homogeneity of variance were assessed with Shapiro-Wilks and Barlett tests, respectively (Sokal and Rohlf, 1995). *A posteriori* comparisons for the rmANOVA models between experimental groups were performed by the comparison of their Estimated Marginal Means (EMMs) by factor levels as supported by the function pairs in the *emmeans* R-package (Lenth, 2018). The Kaplan-Meier survival functions were derived from the censored data of each experimental group using the survfit function from the *survival* R-package (Therneau, 2015). To determine the effect of the experimental factors (i.e.: endophyte infection and saline stress) on the survival probabilities, a Cox proportional-hazard model was performed for each species data using the coxph function of the same package. Further pairwise comparisons were performed using the Peto and Peto modification of the Gehan-Wilcoxon test as allowed in the pairwise_survdiff function, implemented in the *survminer* R-package (Kassambara & Kosinski, 2018). The assumption of proportionality between experimental factors for proportional hazard models was verified with the cox.zph R-function (Therneau, 2015).

## RESULTS

### Crop yield

While the final yield of lettuce and tomato crops was reduced by plus 150 mM NaCl solution saline stress, endophyte inoculation increased final yield (Fig. 1A). The interaction factor (E × S) revealed that the impact of endophyte inoculation on final yield was stronger under saline stress both in lettuce (*F*_1,96_ = 387.7; *P* < 0.0001) and tomato (*F*_1,96_ = 139.2; *P* = < 0.0001).

**Figure 1.**
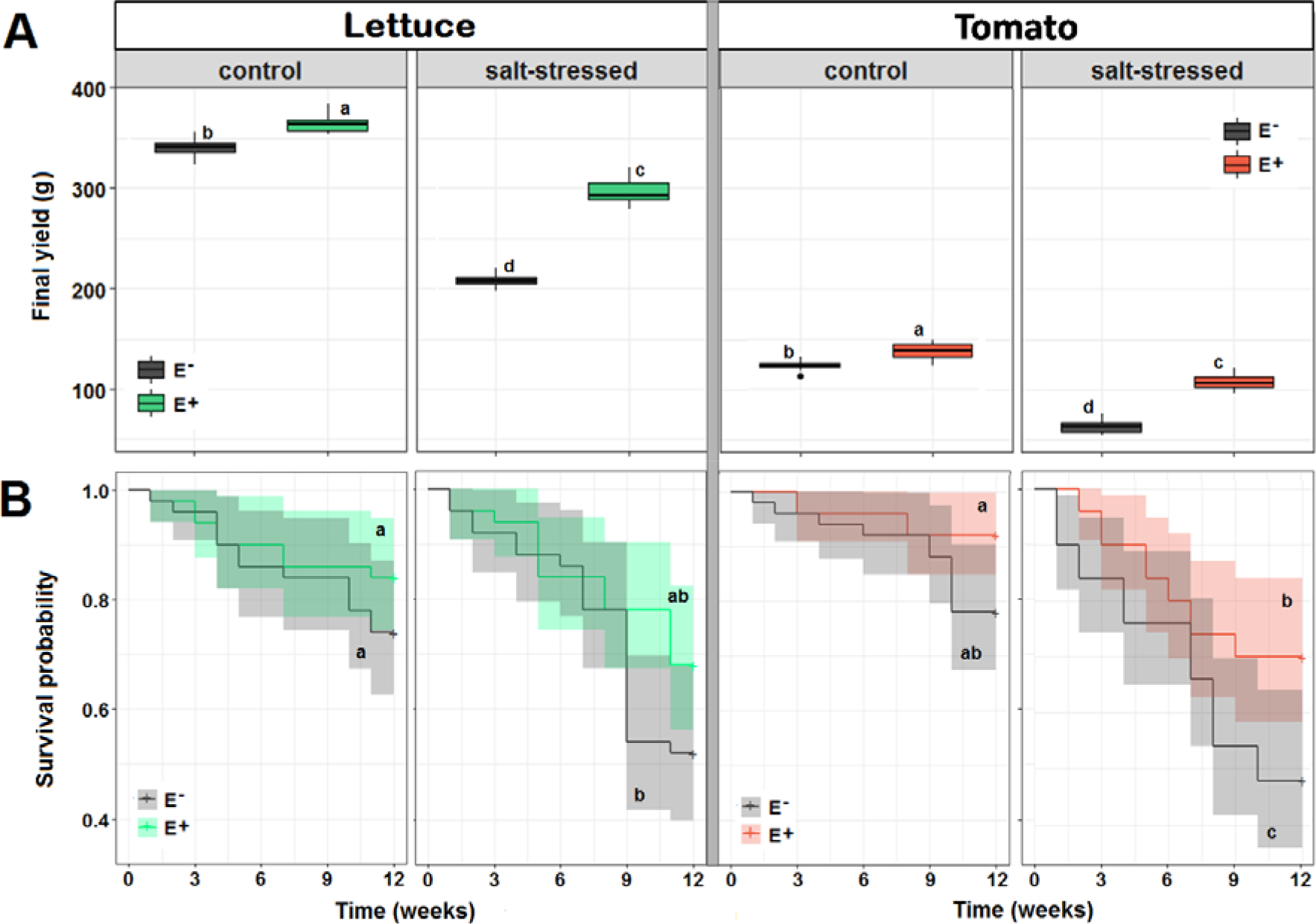
Effects of saline treatment (control vs. salt-stressed) and fungal inoculation (E− vs. E+ plants) on: (A) yield at end of experiment and (B) survival probability along time, in lettuce and tomato plants along time. Different lowercase letters indicate significant differences between treatments (Tukey HSD tests; α = 0.05) independently for each species. The box and bars in the plot represents the interquartile distribution of the date in each experimental group.

The survival probabilities of both species appeared to be significantly affected by the presence of the endophyte, as well as by the saline stress (Cox lettuce model: *n* = 200, mortality events = 139; *df* = 3; likelihood ratio = 11.8, *p* = 0.0027; Cox tomato model: *n* = 200, mortality events = 144; *df* = 3; likelihood ratio = 26.7, *p* = < 0.0001). Specifically, the Cox model suggest that in general for lettuce the absence of the endophyte results in a 1.6 fold hazard increase, while growth under saline stress leads to a higher risk of dead increased by a factor of 2.1 (Sup. Table 1). On the other hand, despite similar, in tomato both factors were more relevant for survival; non-inoculated plants were 2.1 times more prone to die if compared with those with the endophyte consortium, whereas the saline stress increased the hazard in 3.4 times (Sup. Table 1). Interestingly, despite that saline stress seems to drive the main survival differences in both species, the presence of the endophyte significantly reduces the individual mortality in both crops under stress (Fig. 1B). This effect is particularly observed on tomato plants, which survival probabilities felt below 50% under saline stress when the endophytes were not added.

### Ecophysiological Traits

Salt stress and fungal inoculation significantly affected A_max_ and WUE in both crops (Table 1). Moreover, saline stress reduced *A*_*max*_ in time; in contrast, endophyte inoculation increased A_max_ along the experiment (Fig. 2A, Table 1). The lack of interaction between endophyte inoculation and saline stress indicates that the increase in *A*_*max*_ due to endophytes was not affected by salinity. Saline stress reduced WUE in both species only among not inoculated individuals, while E+ plants increased their WUE levels along time (Fig. 2B, Table 1). Without saline stress (control), the positive effect of the endophyte on WUE was significant in lettuce but not in tomato plants, despite the significant changes in time observed for both crops (Fig. 2B). This was consistent with the significant interaction between endophyte inoculation and saline stress (Table 1). Notably, under saline stress endophytes increased WUE exceeding the maximal values of controls in both species.

**Figure 2.**
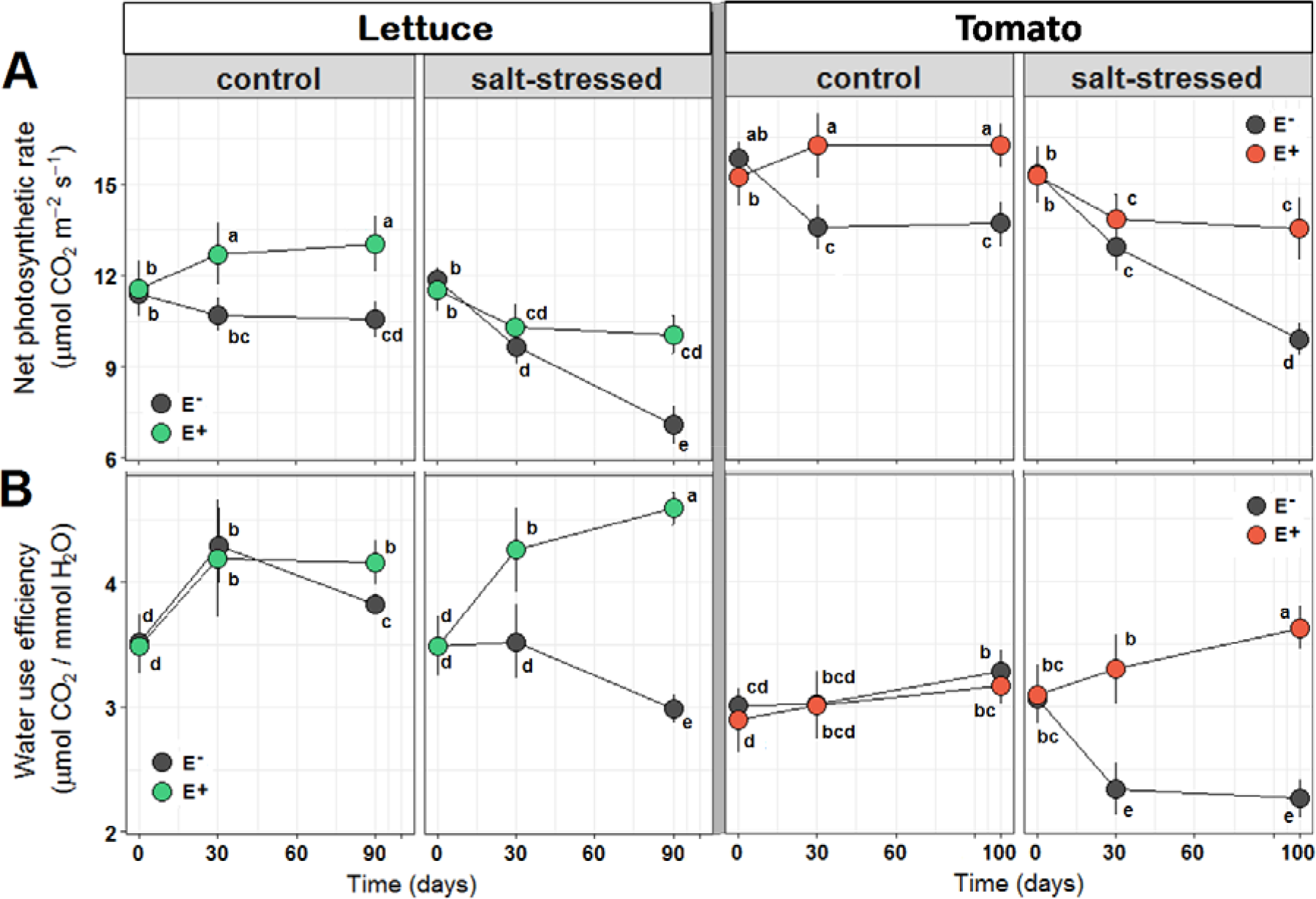
Effects of saline treatment (control vs. salt-stressed) and fungal inoculation (E− vs. E+ plants) on: (A) the net photosynthetic rate (*A*_*max*_) and (B) water use efficiency (WUE) of lettuce and tomato plants along time. Different lowercase letters indicate significant differences between treatments (Tukey HSD tests; α = 0.05) independently for each species. Circles are means (± 1SE).

**Table 1.**
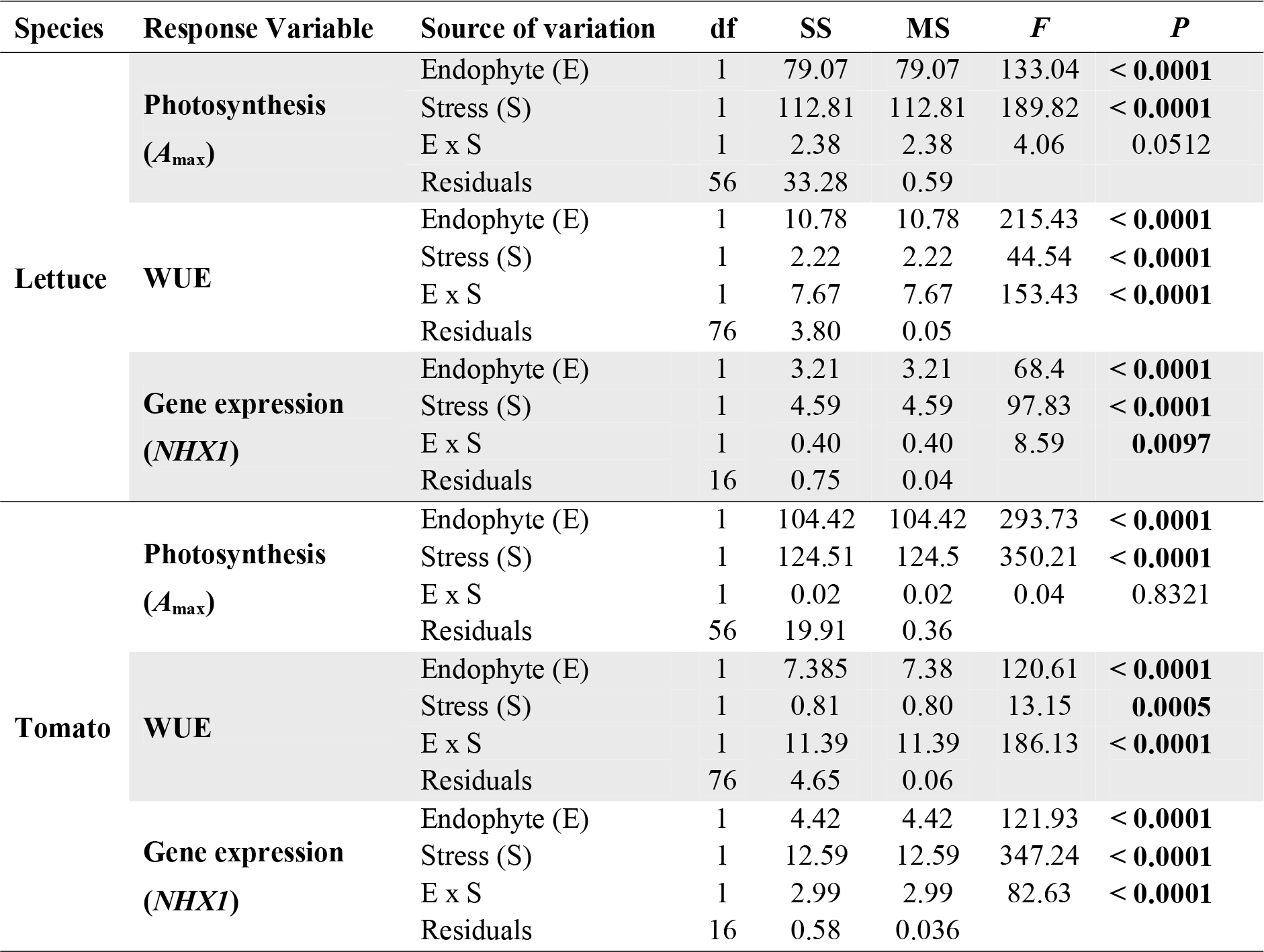
Results of repeated measures ANOVA’s assessing the effects endophyte inoculation (E) and saline stress (S) on the net photosynthetic rate (*A*_*max*_), water use efficiency (WUE) and *NHX1* gene fold expression in lettuce and tomato plants. Significant probability values (*p* < 0.05) are highlighted in red. The results for the factor “time” and its interactions with E and S are not included since all of them resulted highly significant (*p* < 0.001) for both species in the three evaluated traits.

### Gene Expression and content of foliar Na^+^

The relative expression of the *NHX1* gene showed a significant increase in time as a result of both, saline stress and endophyte inoculation (Fig. 3A, Table 1). However, despite significant changes in time were observed under control conditions in both crops, only in lettuce the relative expression of *NHX1* gene was increased by fungal endophytes (Fig. 3A). The significant interaction between those factors indicates that, under saline stress, endophyte inoculation induced a higher increase in the expression of *NHX1* gene. In both species the maximal gene expression was reached after 30 days (Fig. 3A).

**Figure 3.**
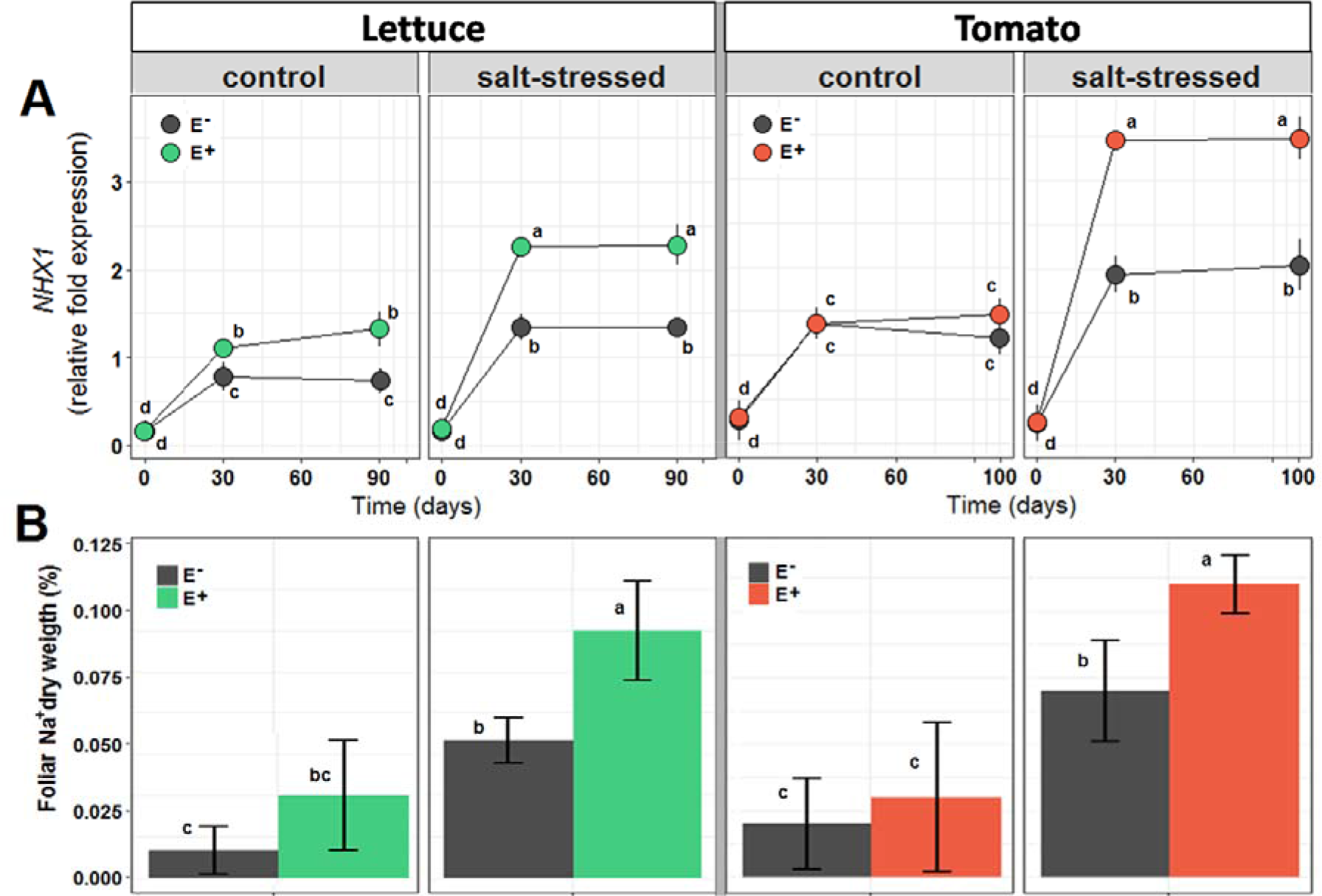
Effects of saline treatment (control vs. salt-stressed) and fungal inoculation (E− vs. E+ plants) on: (A) the expression of *NHX1* gene fold and (B) foliar Na^+^ content of lettuce and tomato plants along time. Different lowercase letters indicate significant differences between treatments (Tukey HSD tests; α = 0.05) independently for each species. Circles and bars are means (± 1SE).

The percentage of foliar Na^+^ was significantly affected by saline stress and endophyte inoculation (Fig. 3B). For both species, lettuce and tomato, there were no differences between E− and E+ plants under normal salt availability. In contrast, under salt stress E+ plants from both species accumulated a significantly higher proportion of Na^+^ than E− plants (Fig 3B).

### Nutrient content

In lettuce, the percentage of nitrogen and phosphorus content were significantly increased due to endophyte inoculation both under control and salt stressed conditions (Table 2). Besides, plants inoculated with endophytes grown under salt stress reached the highest foliar potassium concentration (Table 2). In tomato’s, endophytes increased nitrogen content under control and saline stress. In contrast, phosphorus concentration was greater under control conditions than under saline stress, but was not affected by the endophyte inoculation (Table 2). Potassium concentration, in turn, was higher under saline stress than under control conditions but was not affected by endophytes. Under control conditions, endophytes significantly increased potassium availability (Table 2).

**Table 2.** Concentrations of nutrient (%) present in foliar tissues of lettuce and tomato plants grown under two contrasting levels of salt: control (50 mM NaCl) and salt-stressed (150 mM NaCl), and two inoculation conditions: with (E+) and without root fungal endophytes (E−). Values are means (±SD). Different lowercase letters indicate significant differences between treatments within each nutrient (Tukey HSD tests; α = 0.05).

| Species | Nutrient (%) | Control |  | Saline stress |  |
| --- | --- | --- | --- | --- | --- |
|  |  | E+ | E– | E+ | E– |
| Lettuce | Nitrogen | 3.5a (± 0.16) | 3.1b (± 0.13) | 2.8c (± 0.18) | 2.2d (± 0.22) |
| | Phosphorus | 0.7a (± 0.11) | 0.6a (± 0.09) | 0.4b (± 0.11) | 0.3b (± 0.11) |
| | Potassium | 7.1b (± 0.81) | 6.9b (± 1.07) | 10.1a (± 0.54) | 8.2b (± 0.89) |
| Tomato | Nitrogen | 6.1a (± 0.14) | 5.4b (± 0.17) | 3.6c (± 0.21) | 2.9d (± 0.19) |
| | Phosphorus | 0.8a (± 0.11) | 0.7a (± 0.12) | 0.3b (± 0.11) | 0.3b (± 0.09) |
| | Potassium | 9.5b (± 0.33) | 7.8c (± 0.42) | 12.7a (± 0.39) | 11.5a (± 0.41) |

## DISCUSSION

The combination of growing food demand, inappropriate agricultural practices and global climate change has promoted the rapid expansion of saline soils worldwide, particularly among arid and semiarid environments (Singh et al., 2015). Under such conditions, the physiological and biochemical functions of both wild and cultivated plant species are impaired causing severe yield losses (Deinlein et al., 2014). Therefore, alternative and strategic tools to improve agricultural production are needed for the upcoming and challenging years (Roy et al., 2014). In this direction, we experimentally tested the potential for Antarctic root-fungal endophytes, *Penicillium chrysogenum* and *Penicillium brevicompactum*, to improve the saline-stress tolerance in non-natural (i.e. new) hosts; in this case, two commercial crops, lettuce and tomato. In support for our hypothesis, our results showed the great potential of the established functional symbioses in the two tested crop species by linking the presence of the Antarctic fungal endophytes with positive changes at different biological levels, from gene expression to eco-physiological plant performance. Specifically, the benefits of inoculating the roots of plants with the Antarctic fungal endophytes were more evident under stressful conditions. Endophyte-inoculated plants of lettuce and tomato showed an enhanced survival and yield under salt stress conditions, probably as a result of an improved maximal photosynthesis and water use efficiency. We also determined that the presence of the root fungal symbionts affected the gene expression of both host plants under saline soils. In particular, we observed an up-regulation of the well-known sodium transporter *NHX1* gene in endophyte-inoculated plants that also presented higher concentrations of foliar Na^+^. This latter evidence directly links the root endophytes presence with a functional mechanism to ameliorate saline stress in crops.

Due to their ability to promote plant-growth and stress tolerance, extremophile microorganisms – fungi and bacteria – have received considerable attention in the last few years. The Antarctic ecosystems are among the harshest environments worldwide. Together with cold and arid environmental conditions, Antarctic vascular plants are challenged by saline conditions, as they are restricted to ice-free coastal zones near seashores. Thus, this particular combination of environmental stresses is expected to favors symbiotic associations between soil microorganisms and Antarctic vascular plants though natural selection. A growing number of studies have shown that plant-microorganism’s mutualisms are common at extreme environments such as deserts (Oelmüller et al., 2009; Sangamesh et al., 2018), high-mountain (Molina-Montenegro et al., 2015; Kotilinèk et al., 2017) or polar ecosystems (Upson et al., 2009b; Newsham, 2011a; Torres-Díaz et al., 2016), for which those microbial symbionts appear to be great candidates for biotechnological applications (Khan et al., 2013; Rho et al., 2017).

The number of soil fungi that are reported as beneficial for host plants by establishing endophytic colonization of the roots is increasing (Harman et al., 2004; Rodriguez et al., 2009). Several studies have found evidence showing that fungal endophytes isolated from semiarid environments, such as *Penicillium janthinellum* (Khan et al., 2011a), *Penicillium funiculosum* (Khan et al., 2011b), or *Paelomyces formosus (*Khan et al., 2012), can improve plant growth under salt stress through secretions of phytohormones alleviating the effects of salt stress. Different strains of *Trichoderma*, likely one of the most studied soil fungi, are known to deliver benefits on a number of host species through boosting the physiology and growth of plants (Harman et al., 2004; Harman, 2011). The two Antarctic fungal endophytes used here, *Penicillium chrysogenum* and *Penicillium brevicompactum*, had previously been found responsible of high lettuce yield under water shortage conditions (Molina-Montenegro et al., 2016a). As they did also here, those endophytes improved plant physiological parameters (i.e., *A*_*max*_ and WUE) and the accumulation of the amino acid proline (Molina-Montenegro et al., 2016a), an osmotic molecule that has been previously associated to other symbionts conferring plant tolerance to water deficit (Nagabhyru et al., 2013; Ortiz et al., 2015). Interestingly, the beneficial effects of these Antarctic fungal endophytes seems to be ubiquitous as they are also able to establish functional symbioses with plant species from phylogenetically unrelated plant families such as the native Antarctic vascular plant *Colobanthus quitensis* (Torres-Díaz et al., 2016), three endangered shrubs from semiarid environments *Flourensia thurifera* (Asteraceae), *Puya berteroniana* (Bromeliacae) and *Senna cumingii* (Fabacae) (Molina-Montenegro et al., 2016b; Fardella et al., 2014) and the endangered south American tree *Nothofagus alessandrii* (Torres-Díaz, unpublished results).

The different benefits plants obtain from symbiotic associations with fungal endophytes are well established; but the identification of the underlying mechanisms to those effects can be elusive (Rodriguez et al., 2009; Hamilton et al., 2012). Apart from being source of bioactive secondary metabolites that provide crop resistance to abiotic stress, endophytic fungi can modulate the entire expression of the genome getting the plant ready to tolerate or defense (Harman et al., 2004; Hamilton et al., 2012; Khan et al., 2013, 2015). For example, ABA–deficient tomato mutant plants (*Sitiens* mutants) grown in association with the endophyte *P. janthinellum* LK5 resisted salt stress by producing gibberellins and activating defensive mechanisms in the host *Sitiens* plants (Kahn et al., 2013). Similarly, culture filtrates of the endophytic fungus *Paelomyces formosus* LHL10, extracted form cucumber plants, increased the growth of mutant rice *Waito*-C cultivar (gibberellins deficient) and cucumber shoots under salinity conditions (Khan et al., 2015). The authors found that *P. formosus* produced the increased gibberellins and the auxin IAA (Khan et al., 2015). Besides, the cucumber plants associated to *P. formosus* counteracted saline stress through accumulation of proline and antioxidants, being able to maintaining water potential and membrane functionality (Khan et al., 2015).

In our work, a remarkable observation for both crops under salt stress was the link between the induction of an enhanced *NHX1* transcription and the greater foliar ion accumulation (K^+^ and Na^+^ on lettuce, and Na^+^ on tomato) among endophyte-inoculated plants. In relation with abiotic stress, different *NHX* antiporters have been related with the regulation of diverse factors including cell expansion (Apse et al., 2003), internal pH (Leidi et al., 2010), root-growth (McCubbin et al., 2014) and ionic homeostasis (Bassil and Blumwald, 2014). Regarding salt tolerance, the ionic homeostasis related with the regulation of the K^+^ and Na^+^ within the cytosol has been denoted as the main function of the *NHX* genes (Hernandez et al., 2009; Bassil and Blumwald, 2014). An accepted mode of action results in the transport of either K^+^ or Na^+^ into cellular vacuoles, controlling the cellular pH and its internal turgor (Bassil and Blumwald, 2014). This would allow the normal cellular functioning even under environments with increased saline concentrations, for which the vacuolar accumulation of this ion is a determinant feature for plants under osmotic stress (Jiang et al., 2010).

Plants can use different mechanisms to cope with saline sodic soils. Together with the regulation of the cellular osmotic balance by the activation of specialized pathways that control the export and compartmentalization of sodium cations, plants can adjust root architecture by for example, guiding growth toward less salty soils (Galvan-Ampudia and Testerink, 2011; Deinlein et al., 2014). Although we did not assessed morphological changes in the root system of plants, the greater relative growth and higher photosynthetic rate ‒by the end of the experiments‒ could be at least partially attributed to the positive effect of endophytes on the root functionality. For example, a significant positive growth seems to be associated to root colonization by *Trichoderma* strains (Harman, 2011). An endophyte-mediated improvement of root systems is also supported by the higher concentration of N in foliar tissues from inoculated plants.

Although highly promising, the applied use of beneficial microorganisms to improve plant protection and performance in agriculture, is still not widely accepted because the output of symbiotic interactions are susceptible to vary (i.e., positive, neutral or negative) in response to host, and/or environmental conditions (Ruotsalinen et al., 2004; Gundel et al., 2013; Murphy et al., 2014; Kauppinen et al., 2016; Billingsley-Tobias et al., 2017; Wei and Jousset, 2017). It is interesting to remark that the two crop species responded to the endophyte inoculation with higher photosynthetic capacities and positive yields even under the control conditions. Thus, our work is in accordance with the results from a recent meta-analysis which shows that the overall effect size to the endophyte inoculation on plant growth is positive in both stressed and not-stressed conditions, showing no host-specificity (Rho et al., 2017). In this sense, the positive responses of plants to endophyte inoculations indicate that the referred symbioses could be used as a biotechnological tool not only for crops but also for plants (wild or cultivated) inhabiting marginal environments or environments that are challenged by the climate change.

## Acknowledgements

We acknowledge the logistic support of the Chilean Antarctic Institute (INACH). M.A. Molina-Montenegro is supported by project FONDECYT 1181034.

## Author Contributions

MAM-M and CT-D designed the experiments. MAM-M, ISA-R and CT-D performed the experiments. MAM-M, ISA-R and PEG analyzed and discussed the data. MAM-M wrote the paper along with IA-R, CT-D and PEG. All authors reviewed the manuscript.

---

**Supplemental Table 1:** Cox proportional-hazards model coefficients for letucce and tomato as derived from the censored deads among experimental individuals during 12 weeks. The presence or absence of the endophyte consortium on the plant host and the saline regimes (control and stressed) in which they growth were analyzed. HR: hazard ratio, SE: standar error, *z*: z statistic, *p*: probability value. Values of *p* under 0.05 are highligted in red and suggest statistical significance.

| Species | Factor | $\beta$ | HR ($e^{\beta}$) | SE | $z$ | $p$ |
| --- | --- | --- | --- | --- | --- | --- |
| Lettuce | Endophyte (absent) | 0.514 | 1.672 | 0.262 | 1.96 | 0.048 |
|  | Saline condition (stressed) | 0.748 | 2.113 | 0.27 | 2.77 | 0.0056 |
| Tomato | Endophyte (absent) | 0.788 | 2.198 | 0.282 | 2.79 | 0.0053 |
|  | Saline condition (stressed) | 1.235 | 3.439 | 0.302 | 4.08 | < 0.001 |

